# Persistence of florpyrauxifen-benzyl in sediments following application to a large oligotrophic lake to control Eurasian watermilfoil

**DOI:** 10.1101/2025.09.23.678083

**Authors:** B. Wiltse, B. Mattes, C. Navitsky, S. Chakraborty, J.C. Stager, E. Buell, K.C. Rose

## Abstract

Florpyrauxifen-benzyl (FPB) is a recently registered arylpicolinate herbicide used to control dicots including Eurasian watermilfoil (EWM). Laboratory studies indicate rapid degradation of FPB in water, but sediment persistence in aquatic systems remains poorly understood, creating uncertainty in ecological risk assessments. We evaluated the environmental fate and transport of FPB and its degradants in water, plants, and sediment following application of the herbicide to two bays of Lake George, NY. FPB was rapidly lost from the water column within 72 hours, consistent with prior studies, but persisted in sediments for at least one year. Sediment cores revealed vertical migration of FPB, which may contribute to frequent non-detects in surface sediments and potentially explain why US EPA was unable to establish sediment half-lives in aquatic systems. Measured concentrations exceeded the 28-day no observable adverse effect concentration (NOAEC) for chironomids for at least a year, highlighting the potential for long-term chronic exposure to benthic organisms. Modeled sorption and dilution scenarios aligned with field observations and indicated that circulation dynamics expanded the affected area beyond the direct treatment zones. These results demonstrate that FPB and its degradants persist longer in sediments than suggested by laboratory studies, raising important questions about long-term ecological risks, effects of repeated applications, and underestimation of sediment persistence. Long-term studies such as this are essential for understanding the fate of these compounds and for guiding informed decisions about future herbicide applications.

## Introduction

Aquatic herbicides are routinely used to manage and control invasive aquatic macrophytes. As a result, some broadleaf aquatic plants, such as Eurasian watermilfoil (EWM, *Myriophyllum spicatum*), have developed tolerance to herbicides such as 2,4-dichlorophenoxyacetic acid (2,4-D) and fluridone.^1–3^ Additionally, there are growing challenges with EWM hybridizing with native milfoils, resulting in reduced efficacy of commonly used aquatic herbicides.^4,5^ Increasingly, natural resource managers are seeking alternative methods to overcome this tolerance, including alternating the types of herbicides applied and in some cases using combinations of herbicides.^6–8^

Florpyrauxifen-benzyl (FPB), marketed under several trade names and commercial formulations, is a novel arylpicolinate aquatic herbicide that gained approval for use in aquatic environments by the United States Environmental Protection Agency (US EPA) in 2017.^9^ FPB was developed in part to overcome issues of invasive aquatic macrophyte tolerance to other herbicides and to present a lower risk profile relative to other chemical treatments. Laboratory, field, and mesocosm studies on FPB highlight its efficacy in treating EWM with minimal impacts on non-target aquatic macrophytes.^9–15^

Laboratory studies indicate that FPB rapidly degrades in water through photodegradation, biodegradation, and hydrolysis.^16,17^ US EPA’s risk assessment for FPB determined half-lives in water and soils but was unable to determine half-lives in aquatic sediments due to variable and limited detection across the two field studies that were assessed. Soil half-lives in EPA’s assessment ranged from 42-55 days. These studies were conducted in systems that are neutral to alkaline, which would result in shorter half-lives.^9^ Thus, these studies may not be representative of the field conditions in lakes that are neutral to slightly acidic.

It is important to understand the long-term environmental fate, transport, and ecological impacts of both herbicides and their degradants, which are often overlooked in regulatory review processes.^18^A study of five lakes in Wisconsin that were treated with FPB found the herbicide in sediments as long as 50 days post-treatment.^16^ The persistence of FPB degradants in the environment is expected to be much longer than the parent compound and these substances can also exhibit toxic properties.^9^ In fact, the persistence of herbicide degradants and their potential ecological impacts are often under-evaluated.^19–22^ Additionally, processes used to determine half-lives of herbicides in water-sediment systems often yield estimates that are not robust or are under estimated.^23,24^

The uncertainty of FPB and its degradants persistence in sediments implies a poorly characterized impact on benthic organisms and more work is needed to assess impacts on chironomids, mollusks, and other benthic organisms.^9,25,26^ The potential for impacts to benthic organisms is particularly acute if FPB and its degradants persist and repeated applications of the herbicide occur. However, due to its relatively recent registration, there are limited data on the long-term environmental fate, transport, and ecological impacts of FPB, especially in aquatic use cases.^16^ Much of the peer-reviewed research on FPB relates to agricultural use. Additional information is available through government reports associated with regulatory registration, though aspects of some supporting studies are not publicly accessible for review. To our knowledge there are no publicly available studies tracking the environmental fate and transport of FPB in aquatic use longer than 180 days.^16^

In this study, we assess the fate and transport of FPB and its degradants in water, plants, and sediments following the application of the herbicide to two bays in Lake George, NY to control Eurasian watermilfoil. This study spans thirteen months post-application, making it the longest assessment of the fate and transport of the herbicide to date.

## Materials and Methods

Florpyrauxifen-benzyl was applied to Blairs Bay and Sheep Meadow Bay in Lake George, NY by contractors for the Lake George Park Commission on June 29^th^, 2024 to control EWM (Figure 1). The Blairs Bay treatment area was 1.6 hectares with a mean depth of 3.2 meters, resulting in 52,300 cubic meters of treated water. A 24.3-hectare dilution zone was identified beyond the treatment area. The Sheep Meadow Bay treatment area was 1.5 hectares with a mean depth of 4.1 meters, resulting in 59,503 cubic meters of treated water. A 16.2-hectare dilution zone was established beyond the Sheep Meadow Bay treatment area. In each treatment area, the target concentration of the herbicide was 7.72 ug/L. A total of 15.9 liters of the herbicide was applied to Blairs Bay and 18.1 liters to Sheep Meadow Bay.^27–30^

**Figure 1.**
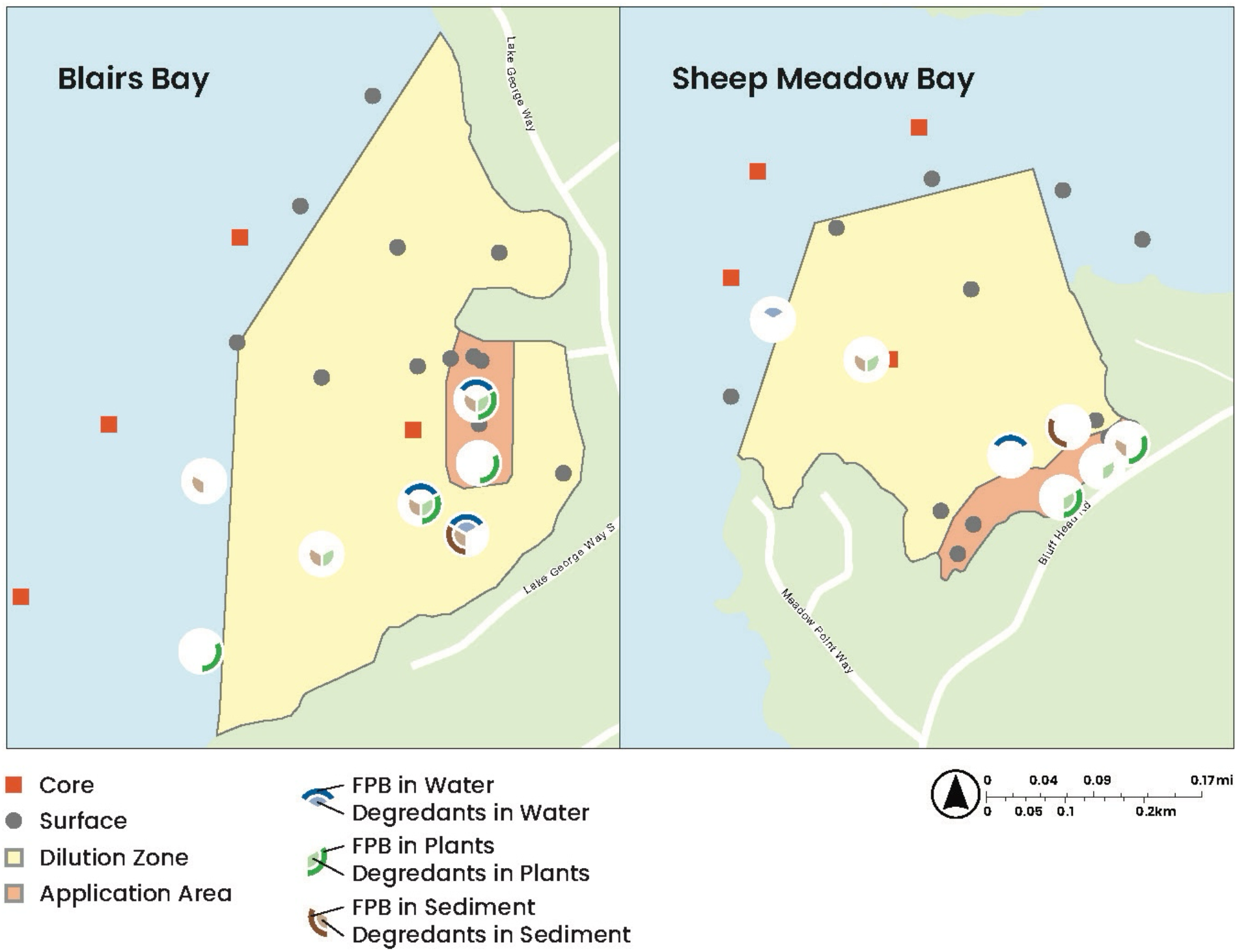
Circles indicate locations where samples were collected over a five-month span in 2024, positive detections of FPB and its degradants from 2024 are shown in the large circles. Red squares indicate locations where sediment cores were collected in 2025, the results from the coring location in the dilution zone are represented in Figure 3.

During and following the herbicide application, water, plant, and sediment samples were collected from each bay. Water samples were collected with a VanDorn sampler at various depths, plant samples were collected by Ponar sampler or hand grabs by divers, and sediments were sampled by Ponar sampler or hand grabs by divers. All samples were stored in glass containers and immediately frozen before transport to the analytical lab.

Samples were collected the day of application, one day later, two days later, one week later, two and a half months later, and five months later. Sampling in the first two days after treatment was most extensive to capture a broad understanding of the immediate fate and transport of the herbicide. Subsequent sampling was targeted to areas where FPB or its degradants had been previously detected to understand long-term fate.

Five sediment cores were collected with a UWITECH gravity core with an internal diameter of 60mm from each bay to better understand the fate of FPB and its degradants in sediments. One core from each bay was collected ten months post-application, and another set of cores was collected twelve months post-application; these cores were taken at the same location within the dilution zones of each bay. Thirteen months post-application, three cores were taken from each bay just outside the dilution zones. Cores ranged from 15 to 20 cm in length and were sectioned at 1 cm intervals into Whirl-Pak bags and immediately frozen before transport to the lab.

All FPB and degradant samples were analyzed at an independent laboratory at the University of Connecticut using ultra high-performance liquid chromatography coupled with a quadrupole time of flight mass spectrometer.

Sediment concentrations were modeled under different sorption (1%, 13%, 25%, 50%, and 100%) and dilution scenarios (treatment area to dilution zone area). We assumed sediment sorption into 1 cm of sediment to calculate theoretical concentrations under these various scenarios.

## Results & Discussion

The detection of FPB and its degradants across each bay was variable spatially, temporally and across different substrates, with most samples showing non-detects (Figure 1 and 2). These results are consistent with other studies assessing the fate of FPB in the environment^9,16,17^ Positive detections for FPB were more common close to the treatment area and immediately after application. It was detected in all three media in both bays following application. Meanwhile, FPB degradants were found in water, plants, and sediments out to the edge of the dilution zones, with one detection of FPB in a plant sample just beyond the dilution zone of Blairs Bay (Figure 1). The spatial pattern of detection, particularly in Blairs Bay, was consistent with lake circulation modeling conducted for the day of treatment.^31^

**Figure 2.**
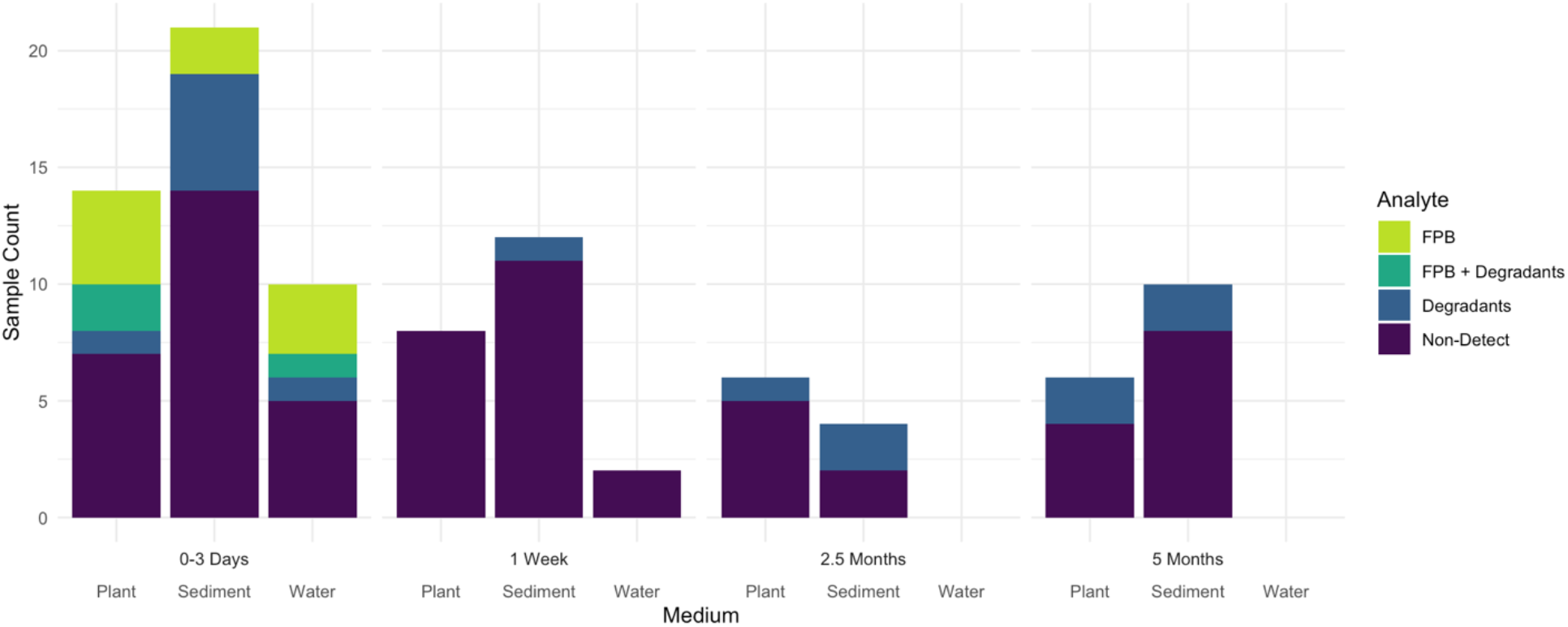
Summary of 2024 sampling for both bays showing detections and non-detects in plants, sediment, and water over a five-month span following the application of FPB.

The transport of FPB and its degradants across the dilution zone highlights the potential for aquatic herbicides, particularly in large lakes with complex circulation patterns, to have effects beyond the targeted treatment area. Our results demonstrate the importance of considering lake circulation when planning herbicide treatments in targeted areas.

We observed FPB and its degradant concentrations drop below the detection limit in water within 72 hours of application, consistent with the literature and manufacturer claims.^9^ However, FPB degradants were also detected in plants and surface sediments five months post-application (Figure 2). Sediment core measurements ten months and one year post application revealed positive FPB and degradant detections in sediments at depths deeper than what would be typically captured by surface sediment sampling (Figure 3). This persistence, which is much longer than previously reported, highlights the limitations of laboratory studies and regulatory review processes that consider limited field studies. In fact, the tendency to underestimate herbicide persistence in sediments in such manner has been routinely documented.^23,24^ The slightly acidic waters of Lake George could potentially contribute to greater persistence of FPB and its degradants. Further studies are necessary to assess photodegradation, biodegradation, and hydrolysis of FPB under neutral to slightly acidic conditions.

**Figure 3.**
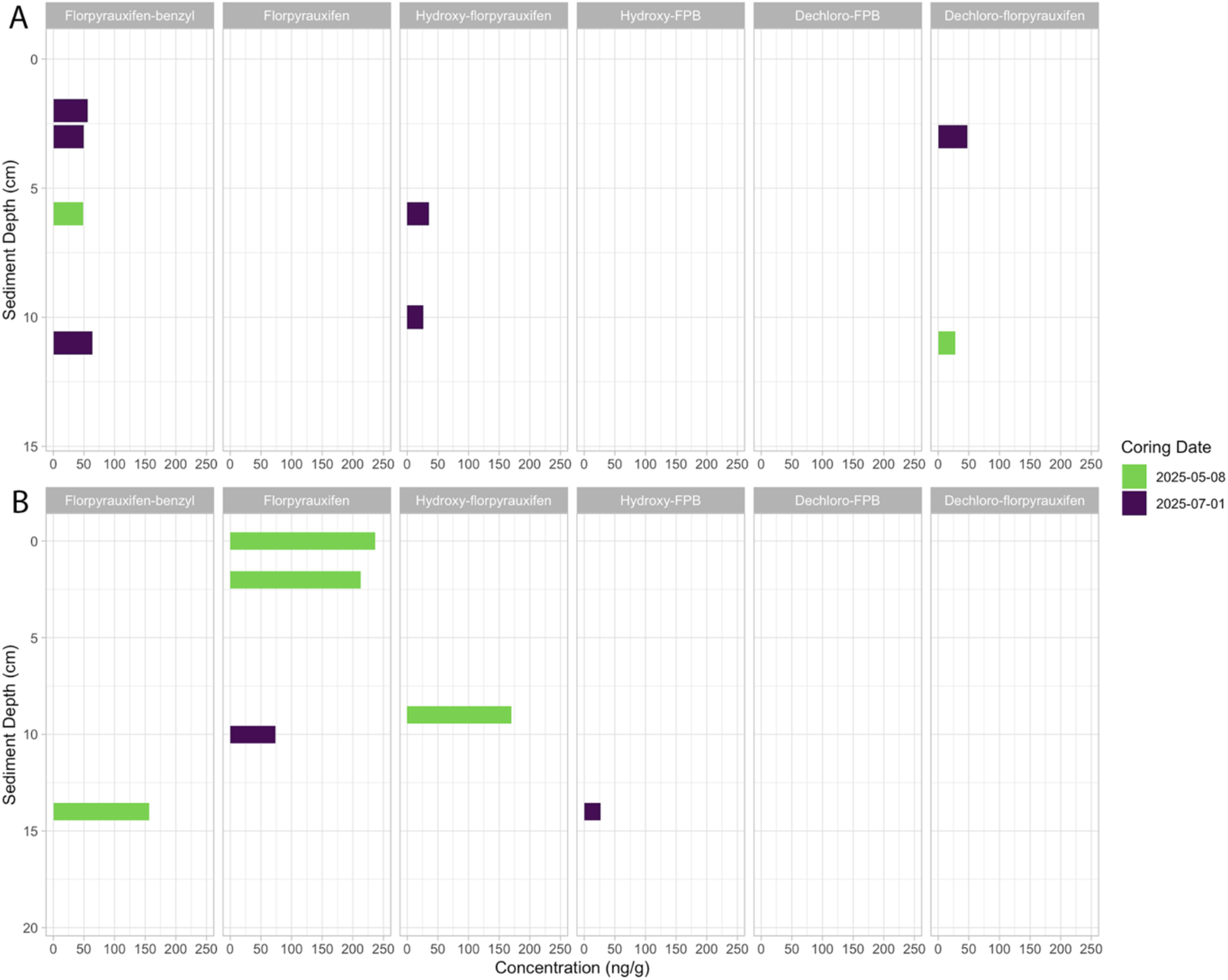
Detection of FPB and its degradants from sediment cores collected from the dilution zone of Blairs Bay (A) and Sheep Meadow Bay (B). Sediment cores were collected at the same location within each bay on the different sampling dates. Results from cores collected from outside the dilution zone are not displayed due to detection of only Hydroxy-FPB at 13cm in one core outside the dilution zone of Blairs Bay.

Sediment core results indicate that FPB moved deeper into sediments than was previously supposed, which may explain the large number of non-detects observed in surface sediment samples. The persistence of FPB at depth in sediments may also explain why the US EPA was unable to determine sediment half-lives for FPB in aquatic use cases.^9^ The US EPA fate and risk assessment indicated low concentrations and variable detection as the reasons for being unable to determine a sediment half-life. EPA relied on two short-term studies of FPB in sediments that may not have looked at sediment profiles. Our findings underscore the importance of comprehensive long-term assessment of the fate and transport of chemical herbicides, points which have been raised elsewhere in the literature.^18,23,24^ Our results also demonstrate that degradation rates in the environment may be far slower than predicted by laboratory studies.

Observed concentrations of FPB and its degradants in surface sediments and sediment cores were above the no observable adverse effect concentration (NOAEC, 25 ng/g FPB) for chironomids identified in the US EPA’s risk assessment.^9^ It is worth noting that the NOAEC identified in the risk assessment was for a 28-day exposure period, whereas our studies indicate persistence at or above these concentrations for at least a year. The impact of long-term exposure of benthic macroinvertebrates to FPB and its degradants is poorly understood.^9,25^ Given the potential for chronic exposure on the order of months to years, as opposed to days to weeks, further research on the impact of FPB and its degradants on these ecologically important organisms is warranted.

To better understand the potential area of persistent exposure to FPB and its degradants in sediments we modelled sediment concentrations under different sorption and dilution scenarios (Figure 4). Broadly, our modeling results are consistent with observed concentrations in the sediment cores, where Sheep Meadow Bay exhibited higher concentrations than Blairs Bay (Figure 3). If we assume sediment sorption rates of 13%, based on previous field studies, we would expect sediment concentrations ranging from 22 to 328 ng/g in Blairs Bay and 32 to 350 ng/g in Sheep Meadow Bay, which are consistent with our observations. Lower observed concentrations in Blairs Bay are consistent with lake circulation modeling that suggested greater circulation in Blairs Bay at the time of treatment and highlights that an herbicide treatment may impact an area several times larger than the direct treatment area.^31^ Notably, our modeling indicates that absent of significant water circulation or dilution, sediment concentrations would be greater than one order of magnitude higher than the 28-day NOAEC for chironomids.^9,16^

**Figure 4.**
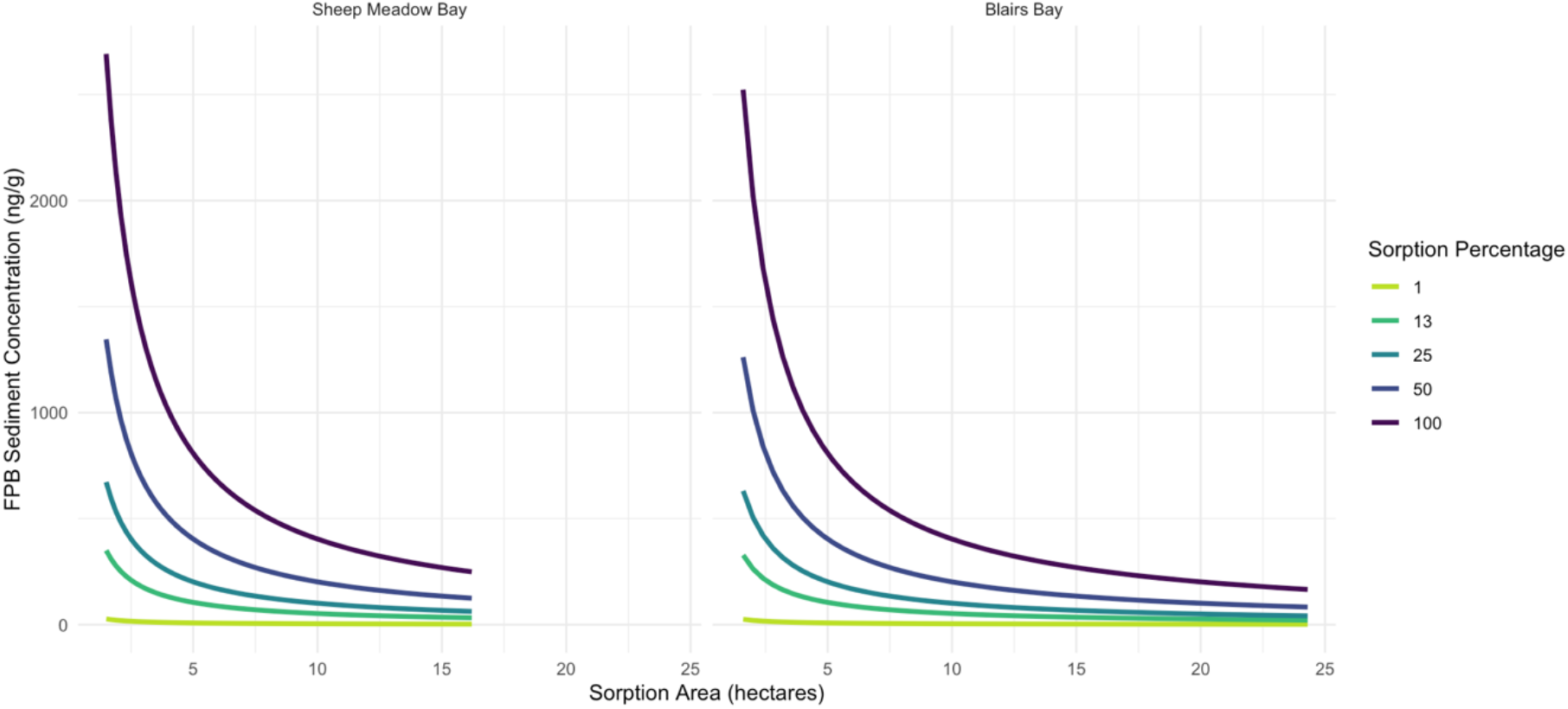
Sediment sorption models showing the potential concentration of FPB in sediments under different sorption rates and across different sediment surface areas. The range of areas for each bay extend from the size of the treatment area to dilution area for each respective bay.

The persistence of FPB and its degradants in the sediments of both bays for much longer than previous studies have reported raises important questions about the potential for long-term chronic exposure to benthic organisms and aquatic macrophytes. Previous studies have shown limited acute effects to mussels but highlight the need to assess sublethal and chronic effects.^25^ The studies that have looked at FPB toxicity often don’t consider long-term chronic effects, the complex interactive effects of exposure to multiple toxic degradants, and the potential for accumulation of these substances in the environment.^9,16,23,24^ Moreover, the root systems of aquatic macrophytes could come into contact with FPB in the sediments, which may explain the slow return of EWM to treatment areas that has been documented in other lakes.^11^ This chronic exposure may eventually contribute to the development of herbicide-resistant strains of EWM.^1,3,5,6^ Other studies have documented hormesis in some species because of low-dose exposure to aquatic herbicides, which could result in increased aquatic macrophyte growth in the areas of treatment.^12,32–37^

Given that our results show FPB persisting in the sediments of both bays at concentrations greater than 50 ng/g and that our study did not attempt to determine sediment half-lives, it is unclear how long FPB could persist in both bays of Lake George. The potential for persistence much longer than anticipated needs to be further studied and has substantial implications for areas that may receive repeated treatments of the herbicide. If FPB persists long enough to last until the next treatment occurs, there is the potential for long-term sediment accumulation.

Furthermore, current literature suggests that the degradants of FPB, which also exhibit toxic properties, can persist for years to decades and therefore are likely to exhibit long-term accumulation in the sediments.^9^ The potential impacts of increasing long-term chronic exposure to these substances because of repeated herbicide application warrants further investigation.

Our findings highlight the importance of evaluating the environmental fate and transport of herbicides in the field over time periods that are typically not captured in regulatory review studies. We have documented FPB persisting in the sediments of Lake George longer than anticipated and shown that it can migrate within the sediment profile. In addition, we have detected FPB and its degradants in water, plants, and sediment across the dilution zones of each treatment area, indicating that the affected area is much larger than the immediate treatment zone. Further research is needed to contextualize these findings, especially in relation to the ecological impacts of long-term chronic exposure to FPB and its degradants. Long-term studies like this are important in determining the fate of these compounds and hence make informed decisions about future applications of the said herbicide.

## Abbreviations

EWM: Eurasian watermilfoil
FPB: Florpyrauxifen-benzyl
NOAEC: No Observable Adverse Effect Concentration
US EPA: United States Environmental Protection Agency

## Acknowledgment

We would like to acknowledge the field staff and scientists at RPI and LGA for their tireless work to sample both bays over arduous and long field days following the application of FPB. We also thank the LGA board of directors for their commitment to the ongoing monitoring and research conducted on Lake George.

## Formula and Equations

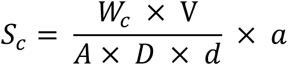

Where;

S_c_ = sediment concentration in ng/g

W_c_ = water concentration in ng/L

V = volume of water treated in L

A = area of sediment exposure in cm^2^

D = depth of sediment sorption

d = density of sediment in g/cm^3^

a = proportion of herbicide sorbed

## References

(1) Richardson, R. J. Aquatic Plant Management and The Impact of Emerging Herbicide Resistance Issues. Weed Technol. 2008, 22 (1), 8–15. 10.1614/wt-07-034.1.

(2) Berger, S. T.; Netherland, M. D.; MacDonald, G. E. Laboratory Documentation of Multiple-Herbicide Tolerance to Fluridone, Norflurazon, and Topramazone in a Hybrid Watermilfoil (Myriophyllum Spicatum M. Sibiricum) Population. Weed Sci. 2015, 63 (1), 235–241. 10.1614/ws-d-14-00085.1.

(3) Busi, R.; Goggin, D. E.; Heap, I. M.; Horak, M. J.; Jugulam, M.; Masters, R. A.; Napier, R. M.; Riar, D. S.; Satchivi, N. M.; Torra, J.; Westra, P.; Wright, T. R. Weed Resistance to Synthetic Auxin Herbicides. Pest Manag. Sci. 2018, 74 (10), 2265–2276. 10.1002/ps.4823.

(4) Moody, M. L.; Les, D. H. Evidence of Hybridity in Invasive Watermilfoil (Myriophyllum) Populations. Proc. Natl. Acad. Sci. 2002, 99 (23), 14867–14871. 10.1073/pnas.172391499.

(5) LaRue, E. A.; Zuellig, M. P.; Netherland, M. D.; Heilman, M. A.; Thum, R. A. Hybrid Watermilfoil Lineages Are More Invasive and Less Sensitive to a Commonly Used Herbicide than Their Exotic Parent (Eurasian Watermilfoil). Evol. Appl. 2013, 6 (3), 462–471. 10.1111/eva.12027.

(6) Kujawa, E. R.; Frater, P.; Mikulyuk, A.; Barton, M.; Nault, M. E.; Egeren, S. V.; Hauxwell, J. Lessons from a Decade of Lake Management: Effects of Herbicides on Eurasian Watermilfoil and Native Plant Communities. Ecosphere 2017, 8 (4). 10.1002/ecs2.1718.

(7) Madsen, J. D.; Wersal, R. M.; Getsinger, K. D.; Skogerboe, J. G. Combinations of Endothall with 2,4-D and Triclopyr for Eurasian Watermilfoil Control; U.S. Army Engineer Research and Development Center: Vicksburg, MS, 2020; pp 1–10.

(8) Ortiz, M. F.; Nissen, S. J.; Dayan, F. E. Endothall and 2,4-D Activity in Milfoil Hybrid (Myriophyllum Spicatum M. Sibiricum) When Applied Alone and in Combination. Weed Sci. 2024, 72 (4), 346–351. 10.1017/wsc.2024.26.

(9) Meléndez, J.; Vogel, V.; Sappington, K. Environmental Fate and Ecological Effects Risk Assessment for the Registration of the New Herbicide for the Use on Rice and Aquatics Florpyrauxifen-Benzyl; U.S. Environmental Protection Agency: Washington, DC, 2017; pp 1–133. https://www.regulations.gov/document/EPA-HQ-OPP-2016-0560-0011.

(10) Beets, J.; Heilman, M.; Netherland, M. D. Large-Scale Mesocosm Evaluation of Florpyrauxifen-Benzyl, a Novel Arylpicolinate Herbicide, on Eurasian and Hybrid Watermilfoil and Seven Native Submsersed Plants. Journal of Aquatic Plant Management 2019, 57, 49–55.

(11) Cattoor, K.; Londo, A.; Walsh, J.; Lund, K. Evaluation of Florpyrauxifen-Benzyl on Invasive Hybrid Watermilfoil in a Central Minnesota Lake. J. Aquat. Plant Manag. 2022, 60 (1), 16–22. 10.57257/japm-20226016.

(12) Mudge, C.; Sartain, B.; Sperry, B.; Getsinger, K. Efficacy of Florpyrauxifen-Benzyl for Eurasian Watermilfoil Control and Nontarget Illinois Pondweed, Elodea, and Coontail Response. 10.21079/11681/42063.

(13) Mudge, C. R.; Sartain, B. T.; Getsinger, K. D.; Netherland, Michael. D. Efficacy of Florpyrauxifen-Benzyl on Dioecious Hydrilla and Hybrid WaterMilfoil - Concentration and Exposure Time Requirements; US Army Corps of Engineers: Washington, DC, 2021; pp 1–17.

(14) Davidson, A. D. Field Application of Florpyrauxifen-Benzyl to Treat Hybrid Eurasian Watermilfoil: Initial Effects on Native and Invasive Aquatic Vegetation. Manag. Biol. Invasions 2023, 14 (3), 467–476. 10.3391/mbi.2023.14.3.06.

(15) Haug, E. J.; Ahmed, K. A.; Gannon, T. W.; Richardson, R. J. Absorption and Translocation of Florpyrauxifen-Benzyl in Ten Aquatic Plant Species. Weed Sci. 2021, 69 (6), 624–630. 10.1017/wsc.2021.38.

(16) White, A. M.; Frost, S. R. V.; Jauquet, J. M.; Magness, A. M.; McMahon, K. D.; Remucal, C. K. Quantifying the Role of Simultaneous Transformation Pathways in the Fate of the Novel Aquatic Herbicide Florpyrauxifen-Benzyl. Environ. Sci. Technol. 2023, 57 (33), 12421–12430. 10.1021/acs.est.3c03343.

(17) Zhou, R.; Dong, Z.; Wang, L.; Zhou, W.; Zhao, W.; Wu, T.; Chang, H.; Lin, W.; Li, B. Degradation of a New Herbicide Florpyrauxifen-Benzyl in Water: Kinetics, Various Influencing Factors and Its Reaction Mechanisms. Int. J. Mol. Sci. 2023, 24 (13), 10521. 10.3390/ijms241310521.

(18) Fenner, K.; Canonica, S.; Wackett, L. P.; Elsner, M. Evaluating Pesticide Degradation in the Environment: Blind Spots and Emerging Opportunities. Science 2013, 341 (6147), 752–758. 10.1126/science.1236281.

(19) Sinclair, C. J.; Boxall, A. B. A. Assessing the Ecotoxicity of Pesticide Transformation Products. Environ. Sci. Technol. 2003, 37 (20), 4617–4625. 10.1021/es030038m.

(20) Cwiertny, D. M.; Snyder, S. A.; Schlenk, D.; Kolodziej, E. P. Environmental Designer Drugs: When Transformation May Not Eliminate Risk. Environ. Sci. Technol. 2014, 48 (20), 11737–11745. 10.1021/es503425w.

(21) White, A. M.; Nault, M. E.; McMahon, K. D.; Remucal, C. K. Synthesizing Laboratory and Field Experiments to Quantify Dominant Transformation Mechanisms of 2,4-Dichlorophenoxyacetic Acid (2,4-D) in Aquatic Environments. Environ. Sci. Technol. 2022, 56 (15), 10838–10848. 10.1021/acs.est.2c03132.

(22) Escher, B. I.; Fenner, K. Recent Advances in Environmental Risk Assessment of Transformation Products. Environ. Sci. Technol. 2011, 45 (9), 3835–3847. 10.1021/es1030799.

(23) Honti, M.; Fenner, K. Deriving Persistence Indicators from Regulatory Water-Sediment Studies – Opportunities and Limitations in OECD 308 Data. Environ. Sci. Technol. 2015, 49 (10), 5879–5886. 10.1021/acs.est.5b00788.

(24) Frost, S. R. V.; White, A. M.; Jauquet, J. M.; Magness, A. M.; McMahon, K. D.; Remucal, C. K. Laboratory Measurements Underestimate Persistence of the Aquatic Herbicide Fluridone in Lakes. Environ. Sci.: Process. Impacts 2024, 26 (2), 368–379. 10.1039/d3em00537b.

(25) Buczek, S. B.; Archambault, J. M.; Cope, W. G.; Heilman, M. A. Evaluation of Juvenile Freshwater Mussel Sensitivity to Multiple Forms of Florpyrauxifen-Benzyl. Bull. Environ. Contam. Toxicol. 2020, 105 (4), 588–594. 10.1007/s00128-020-02971-1.

(26) Mashudi Tangahu, B.V. Review of Florpyrauxifen - Benzyl Herbicide: Bringing Current Global Knowledge for Environmental Impact Mitigation in the Indonesian Context. Jurnal Serambi Engineering 2024, 9 (2), 8733–8739.

(27) Commission, L. G. P. Blairs Bay APA Permit; Adirondack Park Agency, 2022.

(28) Commission, L. G. P. Blairs Bay DEC Permit; New York State Department of Environmental Conservation, 2022.

(29) Commission, L. G. P. Sheep Meadow Bay APA Permit; Adirondack Park Agency, 2022.

(30) Commission, L. G. P. Sheep Meadow Bay DEC Permit; New York State Department of Environmental Conservation, 2022.

(31) Auger, G. A. R.; Kelly, M. R.; Moriarty, V. W.; Rose, K. C.; Kolar, H. R. Understanding Lake Residence Time Across Spatial and Temporal Scales: A Modeling Analysis of Lake George, New York USA. Water Resour. Res. 2024, 60 (2). 10.1029/2022wr034168.

(32) Belz, R. G.; Duke, S. O. Herbicides and Plant Hormesis. Pest Manag. Sci. 2014, 70 (5), 698–707. 10.1002/ps.3726.

(33) Cedergreen, N. Herbicides Can Stimulate Plant Growth. Weed Res. 2008, 48 (5), 429–438. 10.1111/j.1365-3180.2008.00646.x.

(34) Belgers, J. D. M.; Lieverloo, R. J. V.; Pas, L. J. T. V. der; Brink, P. J. V. den. Effects of the Herbicide 2,4-D on the Growth of Nine Aquatic Macrophytes. Aquat. Bot. 2007, 86 (3), 260– 268. 10.1016/j.aquabot.2006.11.002.

(35) Christopher, S. V.; Bird, K. T. The Effects of Herbicides on Development of Myriophyllum Spicatum L. Cultured In Vitro. J. Environ. Qual. 1992, 21 (2), 203–207. 10.2134/jeq1992.00472425002100020008x.

(36) Peres, L. R. S.; Vechia, J. F. D.; Cruz, C. Hormesis Effect of Herbicides Subdoses on Submerged Macrophytes in Microassay Conditions. Planta Daninha 2017, 35 (0), e017165857. 10.1590/s0100-83582017350100076.

(37) Sebastiano, M.; Messina, S.; Marasco, V.; Costantini, D. Hormesis in Ecotoxicological Studies: A Critical Evolutionary Perspective. Curr. Opin. Toxicol. 2022, 29, 25–30. 10.1016/j.cotox.2022.01.002.

